# Simultaneous generation of osteoblasts and adipocytes by a bivalent differentiation medium from bone marrow stromal/stem cells (BMSCs)

**DOI:** 10.1101/2024.01.05.574292

**Authors:** Dawei Qiu, Dong Liu, Jia Chen, Jingwen Huang, Yong Yang, Wanyi Wei, Ziwei Luo

## Abstract

Bone marrow stromal/stem cells (BMSCs) are primitive and heterogeneous cells that can be differentiated into osteoblasts, adipocytes and other subsets. Their bone-fat lineage commitment is responsible for the homeostasis of bone marrow microenvironment. However, there are little effective methods and evidence to simultaneously evaluate the lineage commitment of BMSCs. Here we provide a mixed osteo-adipogenic (OA) differentiation medium that could enable BMSCs differentiation into osteoblasts and adipocytes in vitro, and establish a method for simultaneous assessment of bone-fat lineage allocation of BMSCs based on the Alizarin red S and Oil red O staining, which have been used for detection of specific mineralized nodules and lipid droplets, respectively. This assay provides a specifically simple but effective and low-cost method to visualize the osteogenesis and adipogenesis of BMSCs, and evaluate the differentiation efficiency of lineage allocation of BMSCs.

## 1. Introduction

Bone marrow stromal/stem cells (BMSCs) are primitive and heterogeneous cells resident in the bone marrow, and can be differentiated into osteoblasts, adipocytes, chondrocytes and other cell types in certain conditions[1-3]. In fact, BMSCs are believed to be the common progenitors for both osteoblasts and adipocytes[4, 5]. Under aging, BMSCs are most likely to differentiate into adipocytes instead of osteoblasts, leading to age-related fat accumulation in bone marrow and senile osteoporosis[2, 6], indicating that the lineage choice of BMSCs may preferentially shift towards into adipo-lineage in vivo microenvironment. Therefore, rejuvenation of aged BMSCs that balance the osteogenic and adipogenic lineage commitment of BMSCs is essential for bone homeostasis. However, there are little effective methods and evidence to simultaneously evaluate the lineage commitment of BMSCs in vitro. So far, the vast majority of studies that induce the osteogenic or adipogenic differentiation of BMSCs are conducted separately. For example, when treated with osteogenic medium, only the osteogenic-bias BMSCs will differentiate into osteoblasts[7-9]. But what happens to the adipogenic-bias subsets is largely unknown under the osteogenic condition. Consequently, it is necessary to develop an effective experiment to analyse the fate changes of BMSCs subsets under the same conditions.

Here we provided a special differentiation medium, which contained equal osteogenic and adipogenic drugs, that could directly induce BMSCs differentiation into osteoblasts and adipocytes. This method would provide great convenience for assessment the lineage choice of BMSCs in future studies.

## 2. Materials and methods

### 2.1. Reagents and drugs

All reagents used for cell culture, including low-glucose Dulbecco’s Modified Eagle’s Media (DMEM; Gibco,11054,020), fetal bovine serum (FBS; Gibco,10099,141C), non-essential amino acids (Gibco, 11140, 050), antibiotics (P/S; Gibco, 15070, 063) and trypsin (Gibco,15400, 054) were purchased from Thermo Fisher Scientific Incorporation. Recombinant basic fibroblasts growth factor (bFGF; 100-18B) and epidermal growth factor (EGF; AF-100-15) were purchased from PeproTech Inc..

All drugs used for in vitro differentiation of hBMSCs, including dexamethasone (Sigma; D4902), β-glycerophosphate (Sigma, G9422), L-ascorbic acid (Sigma, A4544), indomethacin (Sigma, I7378), isobutylmethylxanthine (IBMX; Sigma, I7018) were purchased from Merck Group. Insulin (I8830) was from Beijing Solarbio Science & Technology Co., Ltd (Solarbio Inc.). Bone morphogenetic protein 2 (BMP2) was a kind gift from Dr. Erwei Hao (Guangxi University of Chinese Medicine). For detection of osteogenesis and adipogenesis, Alizarin red S (ARS; G8550) and Oil red O (ORO; O8010) powder were purchased from Solarbio Inc..

### 2.2. Cell culture

Human BMSCs were maintained as described before[9, 10]. Briefly, 5×10 ^5^ cells/ml were plated in a 100 mm diameter cell culture dish containing low-glucose DMEM supplemented with 10% FBS, 100 U/ml P/S, 0.1 mM non-essential amino acids, 20 ng/ml bFGF and 20ng/ml EGF. Cells were maintained in a humidified incubator at 37 °C with 5% CO_2_. When 70% to 80% confluent, adherent cells were trypsinized with 0.05% trypsin-1mM EDTA at 37°C for 2 minutes, harvested, and expanded for future use.

### 2.3. Preparation and detection of osteogenesis and adipogenesis

The single and bivalent differentiation of BMSCs was prepared as previously described before[11]. Briefly, 2×10 ^5^ cells per well were seeded into a 12-well plate and then incubated with osteogenic medium, while 1×10 ^6^ cells per well were treated with adipogenic or OA medium. 7 days later, cell cultures were washed with PBS and collected for total RNA isolation. 10 days later, cell cultures were fixed and stained with ARS or ORO to visualize mineralized nodules and lipid droplets[11].

### 2.4. Quantitative real-time PCR

Gene analysis of runt-related transcription factor 2 (*Runx2*) and peroxisome proliferator-activated receptor γ (*Pparγ*) by real-time PCR as we did before[12] were used to validate the bivalent differentiation. Briefly, total cellular RNAs were isolated using RNA Extraction Kits (Solarbio, R1200) according to the manufacturer’s instructions and quantified by Nanodrop One C (Thermo Fisher Scientific, Madison, USA). 1 μg RNA was analyzed by using a TaqMan One Step RT-qPCR Kit (Solarbio, T2210) on LightCycle 96 Instrument (Roche, Mannhein, Germany) according to the manufacturer’s instructions. The primers used for gene amplification were listed as following: for *Runx2*: Forward-5′-TGACATCCCCATCCATCCAC-3′; Reverse-5′-AGAAGTCAGAGGTGGCAGTG-3′; for *Pparγ*: Forward-5′-TATCACTGGAGATCTCCGCCAACAGC-3′; Reverse-5′-GTCACGTTCTGACAGGACTGTGTGAC-3′; for glyceraldehyde phosphate dehydrogenase (*Gapdh*): Forward-5′-ACTCCACTCACGGCAAATTC-3′; Reverse-5′-TCTCCATGGTGGTGAAGACA-3′. The relative expression of target genes was normalized to *Gapdh* and calculated by the 2^−ΔCT^ method.

### 2.5. Statistical analysis

Results are represented as means ± standard deviations. Statistical analysis was performed using Student’s t-test as well as one-way analysis of variance (ANOVA) followed by the Tukey HSD test for post hoc comparison (Origin 8.0, OriginLab). Difference was considered significant when *P*< 0.05 indicated as *, while more significant when *P*< 0.01 indicated as **.

## 3. Results

7 days post induction, mineralized sediments (green arrowheads) and lipid droplets (pink arrowheads) were appeared in their respective wells, but were not evident in the culture dishes treated with OA induction medium (Figure 1A). However, the mineralized sediments and lipid droplets could be clearly distinguished at day 10 (Figure 1B). These results were also confirmed by ARS and ORO staining (Figure 1C, D), suggesting that the OA medium could simultaneously direct osteoblasts and adipocytes formation. By contrast, only osteoblasts were generated in the osteogenic medium, as revealed by ARS and ORO staining and confirmed by gene analysis of *Runx2* and *Pparγ* (Figure 1E, F). The results from adipogenesis were similar to the osteogenesis. Although the lipid droplets stained by ARS were not the same as appeared in ORO staining, the morphological features of adipocytes were preserved intactly, suggesting that adipocytes could be distinguished from osteoblasts by ARS staining (Figure 1D, E, pink arrowheads).

**Figure 1.**
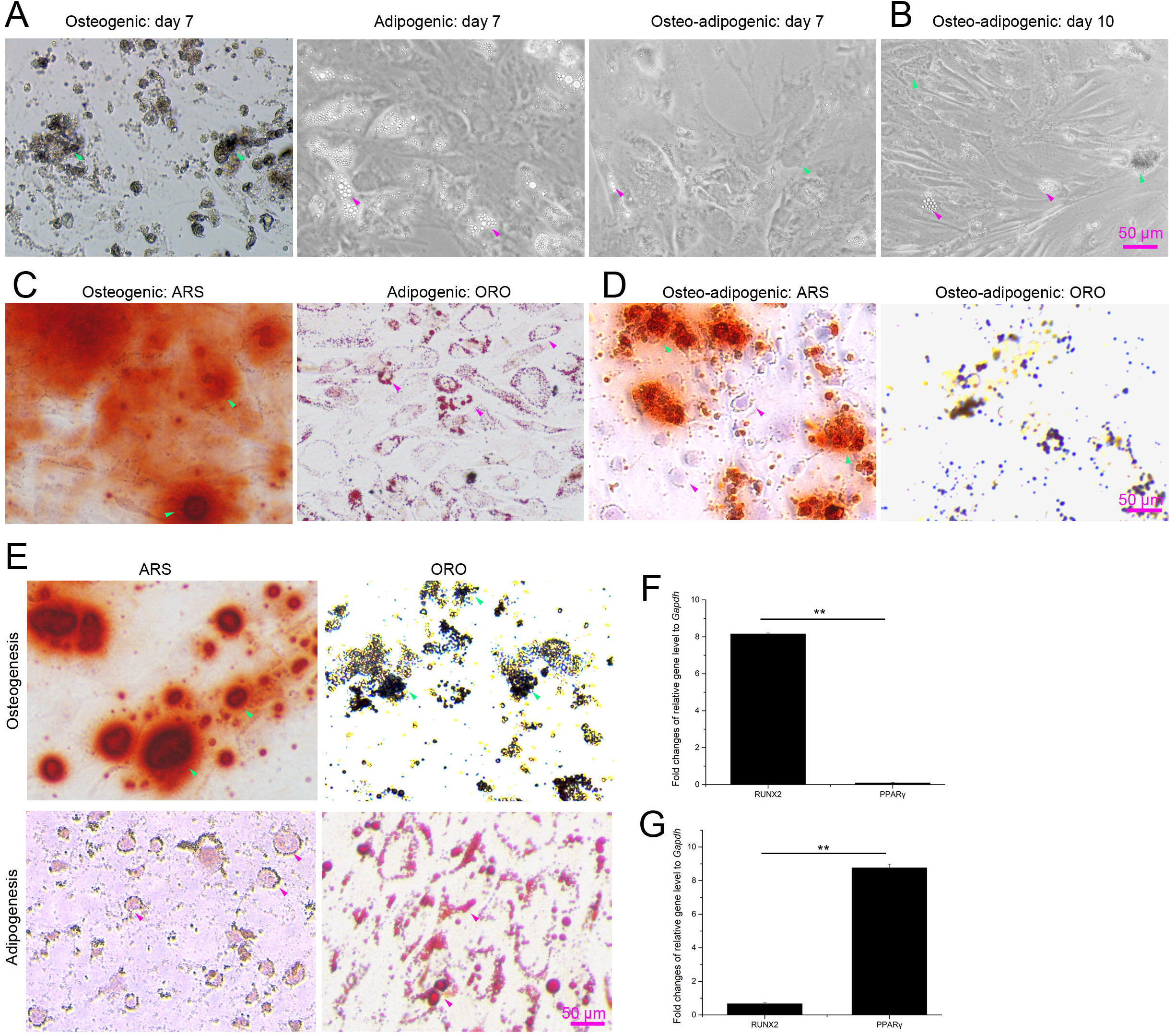
Osteogenic and adipogenic detection of young BMSCs in vitro. **(A, C)** Representative light images (A) and ARS/ORO staining images (C) of cultured BMSCs 7 days post the induction. (**B, D**) Representative light images (B) and ARS/ORO staining images (D) of BMSCs treated by OA induction medium at day 10. (**E-G**) Representative ARS/ORO staining images (E) and gene levels of *Runx2* (F) and *Pparγ* (G) of BMSCs upon to osteogenic or adipogenic differentiation. Results are presented as means ± SD, n ≥ 3. \*\**p* < 0.01.

It is widely accepted that aged BMSCs have decreased abilities to be differentiated into osteoblasts and increased abilities to be adipocytes[1, 13]. In this study, we also used the OA medium to investigate its effects on aged BMSCs. As shown in Fig 2A in the same condition, although aged BMSCs could also be differentiated into osteoblasts and adipocytes, less mineralized nodules (pink arrowheads) and more lipid droplets (green arrowheads) were formed compared with young BMSCs (Figure 2 E and F). In according with the staining, gene level of *Pparγ* was significantly higher than that of *Runx2* in aged BMSCs (Figure 2C). These results validated that aged BMSCs show preferentially differentiation into adipocytes instead of osteoblasts.

**Figure 2.**
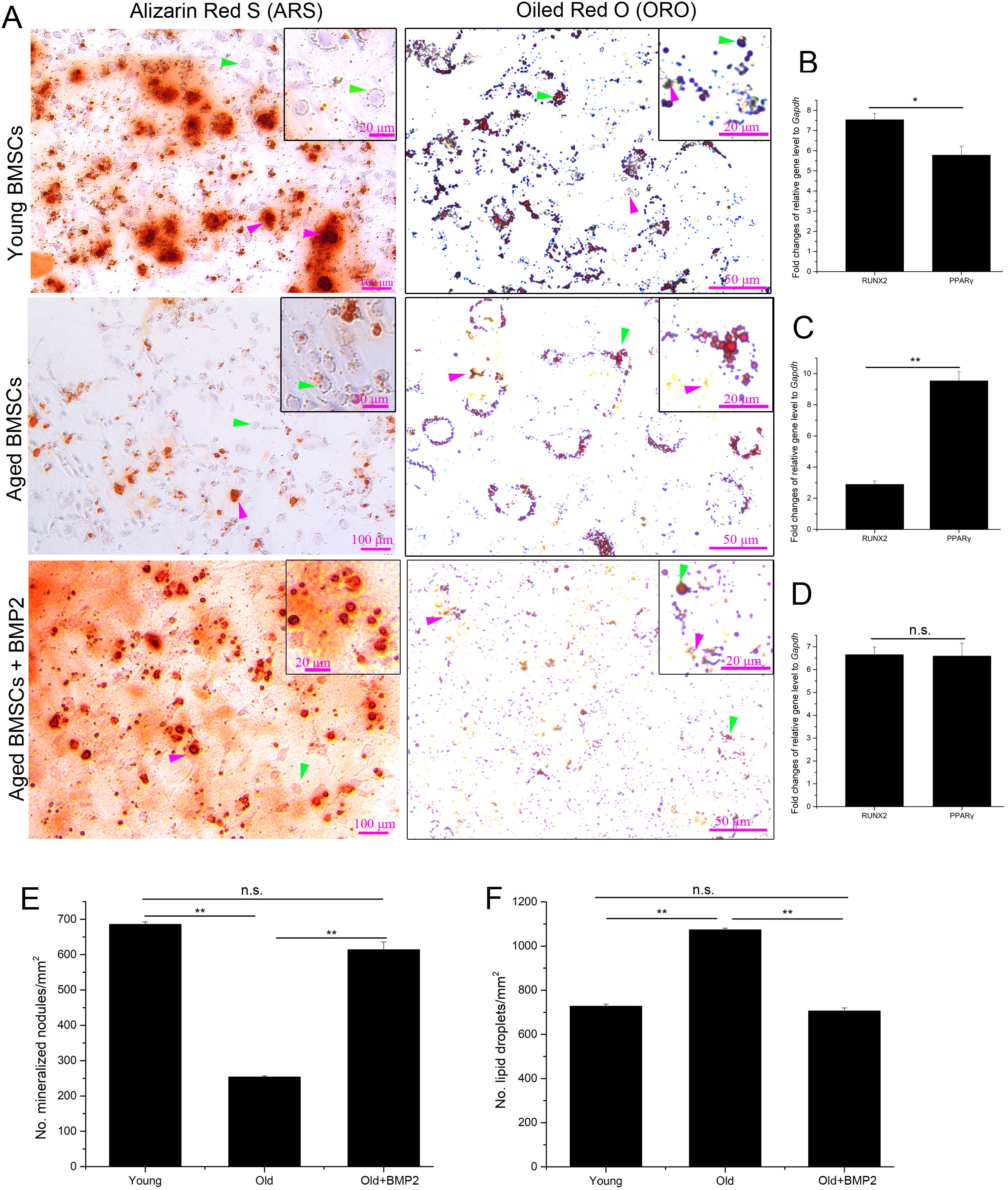
Osteo-adipogenic detection of young and old BMSCs treated with BMP2 or not. **(A)** Representative alizarin red S and Oil red O staining images of young BMSCs (upper), old BMSCs (middle) and BMP2 treated old BMSCs (lower) upon to OA differentiation. Inserts are locally magnified area. **(B-D)** Gene analysis of *Runx2* and *Pparγ* in young BMSCs (B), old BMSCs (C) and old BMSCs treated with BMP2 (D) upon to OA differentiation medium. **(E-F)** Quantitatively analysis of numbers of mineralized nodules and lipid droplets in A. Results are presented as means ± SD, n ≥ 3. \**p* < 0.05, \*\**p* < 0.01, n.s. means no significance.

To better view the fate changes of bone-fat allocation of BMSCs, we additionally treated the aged BMSCs with BMP2, which has been widely employed in many preclinical and clinical studies exploring osteoinductive potential in several animal model defects and in human diseases[14]. Compared with the untreated aged BMSCs, BMP2 remarkedly increased mineralized nodules and decreased lipid droplets under the OA induction (Figure 2A, E, F). It also seemed that BMP2 could restore the differentiation abilities of aged BMSCs (Figure 2E, F), and there was no significance between the gene levels of *Runx2* and *Pparγ* (Figure 2D). These results augmented that it was easy to estimate the fate changes of BMSCs with the help of bidirectional differentiation medium.

## 4. Discussion and Conclusion

Mature cells derived from BMSCs, especially osteoblasts and adipocytes, play key roles in the homeostasis of bone marrow. In turn, as the only tissue where bone and fat coexist in the same microenvironment, bone marrow offers a unique signal to direct the lineage commitment of BMSCs. Therefore, it is reasonably important to uncover the osteogenic and adipogenic choice of BMSCs in the same condition, especially in the study of age-related osteoporosis. To our knowledge, there are little methods to simultaneously determine the osteogenesis and adipogenesis of BMSCs in vitro. In this study, we provided a special differentiation medium and method to monitor the osteoblasts and adipocytes formation of BMSCs. Additionally, our immunofluorescent experiments confirmed the co-exist of RUNX2-positive and PPARγ-positive subpopulations in those BMSCs treated with OA medium in a recent study[15]. This method provided a simple, effective and low-cost but specific approach to directly visualise the lineage choice of osteogenesis and adipogenesis of BMSCs. Moreover, our results not only confirmed that young BMSCs were closer to osteoblasts than adipocytes[5], but also verified that aged BMSCs preferentially generated adipocytes instead of osteoblasts[1, 2, 13]. However, whether the doses of supplemental drugs that were used to induce BMSCs differentiation are optimization combination still needed further studies.

In summary, we have developed a bivalent differentiation medium that permitted both osteogenic and adipogenic differentiation of BMSCs. This discovery would also provide convenience for researching the lineage changes of BMSCs, especially in the case of regulating the skewed differentiation of BMSCs upon drugs against senile osteoporosis, like the research of CXM102 in our work[15].

## Acknowledgements

We sincerely thank Dr. Li Yang (Chongqing University) for the kindly gift of young (14-year-old) and old hBMSCs (68-year-old), and Dr. Erwei Hao (Guangxi University of Chinese Medicine) for the help of cell culture.

## Author Contributions

Conceptualization, Z. Luo, D. Qiu and D. Liu; methodology, Z. Luo, D. Qiu and D. Liu; software, D. Qiu, J. Chen and J. Huang; validation, D. Liu; formal analysis, D. Qiu, D. Liu, Y. Yang and W. Wei; investigation, Z. Luo; data curation, Y. Yang and W. Wei.; drafting the manuscript, D. Liu, W. Wei; reviewing and editing, Z. Luo, W. Wei and D. Liu; visualization, Z. Luo and W. Wei; supervision, Z. Luo, Y. Yang and W. Wei; project administration, Y. Yang; funding acquisition, Z. Luo.

All authors have read and agreed to the published version of the manuscript.

## Data Availability Statement

The data presented in this study are available from the corresponding author, Z. Luo, upon reasonable request.

